# Spatio-temporal dynamics of macroglial cell organisation and proximity to blood vessels during postnatal development

**DOI:** 10.1101/2025.01.14.632893

**Authors:** Naomie Guille, Héloïse Monnet, Tristan Hourcade, Philippe Mailly, Martine Cohen-Salmon, Anne-Cécile Boulay

## Abstract

Brain cortical development results from the proliferation, differentiation, migration and maturation of many cell types. While neuronal development is well characterized, the mechanisms regulating macroglial cells (oligodendrocytes and astrocytes) development remain largely unknown. Recent works suggest that the vascular system plays a key, yet under-evaluated role in this process. In this study, we investigated the spatial organization of macroglial cells within the parenchyma and relative to blood vessels. Using immunolabeling for Sry-box transcription factor (Sox) 9 (macroglial progenitors and astrocytes) and Sox10 (oligodendrocyte lineage), we determined macroglia density, distribution and proximity to blood vessels from postnatal day (P) 1 to P60 in the somatosensory cortex. We showed that Sox9+ cells were evenly distributed across cortical layers with regular intercellular spacing. In contrast, Sox10+ cells concentrated in deeper cortical layers, and exhibited a random distribution. Vascular density and branching increased markedly between P5 and P15 and macroglial cells were closer to blood vessels from P15 onward. We investigated possible alteration of astrocyte distribution in the cortex of MLC1-deficient mice, a model of Megalencephalic leukoencephalopathy with subcortical cysts, in which astrocyte perivascular coverage is altered. No difference with the control condition were found both in young and adult mice, either in the density, distribution or distance to blood vessels. Altogether, we revealed distinct distribution and postnatal development patterns for astrocytes and oligodendrocytes in the brain and in relation to the vasculature.

**Table of Contents Entry:** 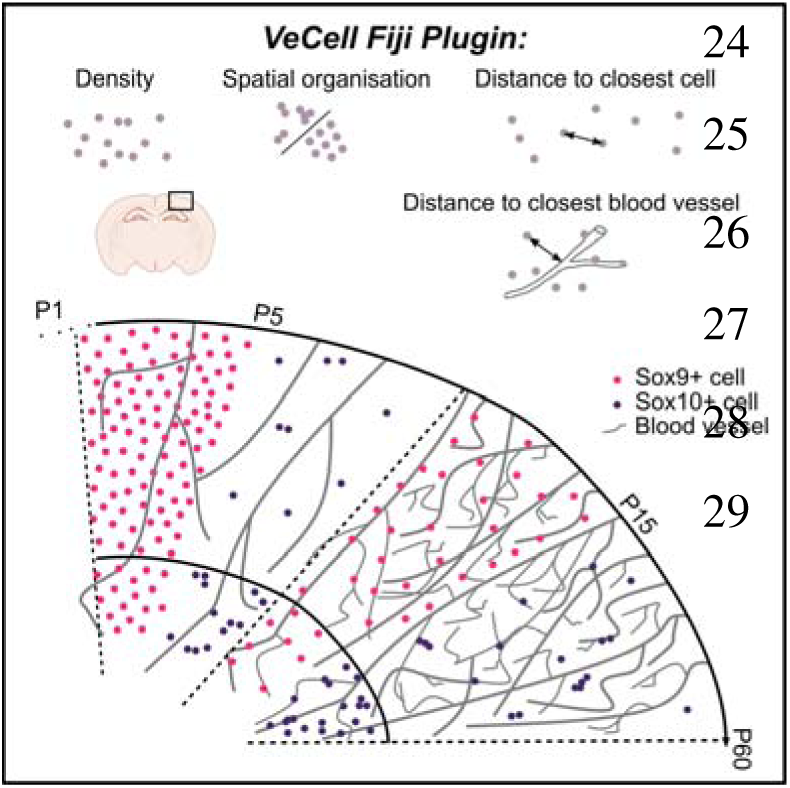

- From P5, Sox9⁺ cells are evenly distributed, unlike Sox10⁺ cells
- From P15, Sox9⁺ cells are located closer to blood vessels than Sox10⁺ cells
- Distribution of astrocytes is not altered in *Mlc1* KO cortex

## Introduction

The mammalian cortex is a complex and highly organized tissue which contains diverse neuronal, glial and vascular cell types. Cortical development is driven by highly regulated and coordinated sequences of cell proliferation, differentiation and migration, and requires the fine tuning of production/elimination and attraction/repulsion mechanisms. Neurons and macroglial cells - astrocytes and oligodendrocytes - are generated from common multipotent neural precursor cells (NPCs) (Miller and Gauthier, 2007). After neuron production, which starts around embryonic day (E) 12 and peaks at E15, NPCs switch to gliogenesis around E18 to sequentially generate astrocyte progenitors and oligodendrocyte precursor cells (OPC). Cortical astrocytes are produced mainly during the first postnatal week from a combination of delaminating NPCs and local proliferation of astrocyte progenitors following colonization of the cortical parenchyma (Clavreul et al., 2019; Ge et al., 2012). From the second postnatal week, their migration is halted and astrocytes mature both molecularly and morphologically to finally tile the brain with non-overlapping domains (Bushong et al., 2004, 2002; Halassa et al., 2007). Oligodendrocyte lineage cells (OligoLC) comprise both oligodendrocyte precursors (OPCs) and myelinating oligodendrocytes (Foerster et al., 2024). During cortical development, OPCs are produced in three spatiotemporal waves. In the postnatal cortex, OPCs generated during the first embryonic wave undergo massive cell death. They are eliminated from the cerebral cortex by P10 and progressively replaced by OPCs locally-produced from cortical progenitors (Kessaris et al., 2006). After birth, OPCs progressively differentiate into oligodendrocytes (OLs) that form the myelin sheath around axons, mostly before the end of the 3^rd^ postnatal week (Orduz et al., 2019), some NG2^+^ OPCs remaining in the mature brain.

Brain vascular network formation is initiated earlier than macroglial cells. Around embryonic day (E)9, the primary vascular network starts to form. From a perineural plexus, vascular sprouts invade the neuroectoderm by vasculogenesis and angiogenesis (Engelhardt, 2003). Postnatally, high angiogenesis rate results in an extensive vascular expansion between P8 and P12. New blood vessels (BVs) arising from ascending veins give rise to a dense capillary bed (Coelho-Santos and Shih, 2020).

BVs and macroglial cells interact during postnatal development. OPCs from ventral origin use blood vessels to migrate to the dorsal telencephalon during embryonic development (Tsai et al., 2016). Whether this also stands for postnatal cortical OPCs remains unknown. The role of BVs in astrocyte progenitor’s migration is also unknown. A significant part of cortical Aldh1l1-positive cells, which include differentiated astrocytes as well as progenitors, are in contact with BVs at P1, but this population drops at P5, in the middle of the highly proliferative and migratory phase of astrocyte progenitors (Clavreul et al., 2019; Freitas-Andrade et al., 2023). BVs have also been shown as a signalling source for both oligodendrocyte and astrocyte maturation (Paredes et al., 2021). A better characterization of how macroglial cells organize and relate to BV during the postnatal period is however necessary.

In this study, we characterized macroglial cell density, distribution and distance to BVs in the developing and mature brain. Macroglial cells were immunolabelled for Sox 9 and 10 which regulate astrocyte and oligodendrocyte differentiation from glial precursors (Stolt and Wegner, 2010). Sox9 is expressed in NPCs and controls their switch to glial progenitors (Güven et al., 2020; Kang et al., 2012; Stolt, 2003). Its expression decreases during OligoLC specialisation, while increasing in astrocytes (Nagao et al., 2016; Klum et al., 2018). Sox10 is expressed in all OligoLCs, which are, during postnatal development, a mix of mature oligodendrocytes and OPCs originating either from cortical postnatal or ventral embryonic progenitors. In these cells, Sox10 regulates the transcription of myelin-forming genes (Foerster et al., 2024; Stolt et al., 2002).

Focusing on the somatosensory cortex, we found that Sox9+ cell density is homogeneous across cortical layers and peaks at P5, while Sox10+ cell density is maintained during postnatal development and is always higher in the deeper layers of the cortex. Sox9+ cells are distributed regularly within the parenchyma from P5. In contrast, Sox10+ cells are randomly distributed with varying distance between cells. Volume and branching of the cortical vascular network increase between P5 and P15. Sox9+ and Sox10+ cells become closer to BVs from P15. Absence of MLC1, an astrocyte protein enriched in perivascular processes, does not alter astrocyte density and distribution.

## Materials and Methods

### Animal experiments and ethical approval

C57BL/6JRj, aged 5 to 60 days were purchased from Janvier labs (Le Genest-Saint-Isle, France) and kept in pathogen-free conditions. Mlc1 KO mice were provided by Raul Estevez (Barcelona, Spain) and maintained on a C57BL6 genetic background (Hoegg-Beiler et al., 2014). All experiments and techniques complied with (i) the European Directive 2010/63/EU on the protection of animals used for scientific purposes and (ii) the guidelines issued by the French National Animal Care and Use Committee (reference: 2013/118). Animal cohorts were made of males and females.

### Brain sections preparation

Mice were sacrificed by cervical dislocation or decapitation depending on mouse age. Brains were carefully collected and fixed in PBS/PFA 4% overnight at 4°C. After dehydration in 30% sucrose, brains were cut into 40-µm-thick (P5 and P15 *Mlc1* KO), 60 µm-thick (Adult *Mlc1* KO) or 80-µm-thick (all other conditions) sections using a Leitz (1400) cryomicrotome and kept at −20°C in storage solution (PBS/glycerol 30%/ethylene glycol 30%).

### Immunofluorescent staining

Brain sections were rinsed three times 15 min in PBS and incubated in a blocking solution (BS) for 1h at room temperature (RT). BS were PBS/normal goat serum 5%/Triton X-100 0.5% (Sox10) or PBS/Triton X-100 0.5% (Sox9). Sections were incubated with primary antibodies diluted in the same BS at 4°C overnight. After three rinses of 15 min in PBS, the sections were incubated for 2h at RT with secondary antibodies and Hoechst reagent, rinsed in PBS, and mounted in Fluoromount G (Southern Biotech, Birmingham, AL).

### Primary and secondary antibodies

Primary antibodies used were polyclonal goat IgG anti-Sox9 (AF3075 Biotechne/R&D, 1:500), monoclonal mouse/IgG2a anti-Sox10 (66786-1-Ig Proteintech, 1:500), AlexaF647-coupled Polyclonal Goat IgG anti-Pecam-1/Cd31 (AF3628R-MTO, Biotechno, 1:300). Secondary antibodies used were AlexaF555-coupled Goat anti-mouse IgG2a (A-21137 thermo Fisher, 1:1000) and AlexaF555-coupled Donkey anti-Goat IgG (A-21432 thermo Fisher, 1:1000). Nuclei were stained using Hoechst (Thermo fisher 62249, 1:2000).

### Image acquisition

Images were acquired using an Axio zoom.V16 microscope (Carl Zeiss) equipped with a 2.3x/0.57 objective, a Flash 4.0 sCMOS camera (Hamamatsu) and an Apotome 2 module. 3D Images were taken with a magnification of x160 and a step size of 4 microns with a variation of 2 to 4 stacks. Mosaic acquisitions were performed using the Tile Scan function with a 10% overlap between adjacent tiles, with 6 or 8 tiles per image. The acquired images were subsequently reconstructed using the Stitching function within the ZEN software. Spinning disk W1 Z-stack projection images were taken with a magnification of x40 and a step size of 0.3 microns.

### Image analysis

Bins used to draw the region of interest (ROI) were defined based on cell density using nuclear staining Hoechst as presented in Figure S1. Bin 1 corresponds to the upper layers of the cortex (II/III-V), Bin 2 to the deeper layer (VI). Layer 1 of the cortex was excluded because of its different organization and morphology of astrocytes. The analysis was conducted in each ROI using a custom-developed plugin for the Fiji software (*VeCell)* (https://github.com/orion-cirb/VeCell) (Schindelin et al., 2012). This plugin integrates the Bio-Formats (Linkert et al., 2010), CLIJ (Haase et al., 2020) and 3D ImageJ Suite (Ollion et al., 2013) libraries. VeCell consists in three analysis steps.

1. <u>Macroglial cell detection:</u> Sox9+ and Sox10+ cells’ detection was automated using the 2D-stitched version of the Cellpose deep learning-based algorithm and *Fiji* software (Schindelin et al., 2012; Stringer et al., 2021). A custom model was trained through the human-in-the-loop workflow available in the Cellpose GUI. This model was initialized with the ‘cyto2’ pre-trained Cellpose model and fine-tuned using a dataset of 28 images collected at three developmental stages (P5, P15, and P60). Of these, 8 images were reserved for validation. The model was trained using the following hyperparameters: learning rate = 0.1, weight decay = 0.0001, and number of epochs = 500. Training and validation loss curves were monitored to ensure convergence. Model performance was evaluated on a separate test set comprising 6 images (2 per developmental stage). The fine-tuned model achieved a significant improvement in mean average precision (mAP) at an IoU threshold of 0.5, increasing from 0.674 (for the ‘cyto2’ pretrained model) to 0.836. In addition, a separate model was trained specifically for P1 developmental stage images to account for the more elongated cell morphology and a lower signal-to-noise ratio. A dataset of 25 images was used for training, with 5 images reserved for validation. The P1-specific model was initialized from the ‘cyto2’ pretrained weights and fine-tuned using similar hyperparameters: learning rate = 0.1, weight decay = 0.0001, and number of epochs = 1000. Model performance was evaluated on a test set of 6 images, demonstrating a substantial improvement in mAP at an IoU threshold of 0.5: from 0.539 for the ‘cyto2’ pretrained model to 0.778 after fine-tuning. These models were deemed suitable for processing the entire dataset. During inference, the following parameters were used: diameter = 20, flow threshold = 0.4, cell probability threshold = 0.0, and stitching threshold = 0.5. Detected 3D cells were filtered based on volume to reduce false positives. Only cells with a volume between 30 µm³ and 3000 µm³ were retained for downstream analysis (Fig. S1).
2. <u>Blood vessel detection:</u> Images were normalized using the Quantile-Based Normalization algorithm (“‘Quantile Based Normalization’ ImageJ PlugIn,” n.d.) ensuring intensity consistency across the dataset. A 3D median filter (*radius_x_ = 4, radius_y_ = 4, radius_z_ = 1*) was applied to reduce noise while preserving vessel borders. Subsequently, a slice-by-slice 2D Difference of Gaussians filter (σ*_1_ = 4,* σ*_2_ = 8)* was applied to enhance vessel structures. A binary mask was generated using the Triangle automated thresholding method. The process was repeated with a second 2D Difference of Gaussians filter (σ*_1_ = 7,* σ*_2_ = 14)*, producing a second binary mask. These two binary masks were then merged using a maximum point operation to ensure comprehensive vessel detection. Post-processing includes applying a 3D closing filter *(radius_x_ = 8, radius_y_ = 8, radius_z_ = 1)* to fill gaps in vessel structures, followed by a 3D median filter *(radius_x_ = 1, radius_y_ = 1, radius_z_ = 1)* to smooth vessel borders. The resulting binary mask was labelled in 3D to identify individual vessel structures, which are filtered by volume to retain only vessels exceeding 600 µm³. Finally, the binary mask was skeletonized using CLIJ library, and branches shorter than 30 µm were excluded to ensure only meaningful segments were included in subsequent skeleton analysis.
3. <u>Computation and saving of metrics for macroglial cells and blood vessels.</u> For macroglial cells, were calculated: volume, distance to the nearest neighbor, mean and maximum distances to the 10 nearest neighbors, distance to the nearest vessel, and nearest vessel’s diameter. Additionally, spatial statistical analysis was performed to identify clustering or dispersion patterns within the macroglial cell population. This was achieved using the 2D/3D Spatial Analysis Fiji plugin from the 3D ImageJ Suite library, which computes the distance function G. This function represents the cumulative distribution of distances between a typical point in the spatial pattern and its nearest neighbor (Andrey et al., 2010). Shorter distances suggest clustering, while longer distances indicate regular spacing (Fig. 2E). The experimental CDF was compared to the CDF of randomly generated patterns within the same area and with the same number of cells. The spatial distribution index (SDI) is derived from this analysis, serving as a metric to quantify the deviation of the observed spatial distribution from a completely random distribution: uniformly distributed SDIs from 0 to 1 reflects a random distribution of cells. SDIs near 0 suggest clustering. SDIs near 1 indicate regular spacing (Andrey et al., 2010). For BVs, skeleton analysis was conducted using the Analyze Skeleton Fiji plugin (Arganda-Carreras et al., 2010). The following metrics are obtained: total volume, total length, mean branch length, number of branches and junctions, mean diameter and diameter standard deviation.

### Statistical analysis

Data shown on graphs are expressed as means and SD. Graphs and statistical analyses were performed using Graphpad Prism 8.0.2 software. For every data, we first verified that they followed a normal distribution using Shapirow-Wilk test. If not, we used Kruskal-Wallis and Mann-Whitney tests. For blood vessel analysis, 2-way ANOVA followed by Sidak’s multiple comparisons test was used to analyse differences between Bins or Tukey’s multiple comparisons test, to analyse differences between ages. For cells density and distribution Mann-Whitney tests were used. For SDI distribution, comparison to normality was performed using Kolmogorov-Smirnov test, comparison to 1 was performed using a Wilcoxon Signed Rank test. For percentages, a chi-square test was used. Alpha thresholds for hypothesis verification were set at 0.05. Statistical tests are indicated in the Figure legends and in the supplementary table 1.

### Use of Artificial Intelligence Generated Content

ChatGPT (version 5) was used to perform the final editing of this manuscript, including grammar and punctuation corrections, as well as improvements in clarity. It was not used to generate any content or the original version of the text.

## Results

### Cortical density of macroglial cells during postnatal development

To characterize the postnatal development of cortical macroglial cells *in situ,* we immunolabelled brain sections from postnatal day (P) 1, P5, P15 and P60 for the transcription factors Sox9 and Sox10, focusing on the somatosensory cortex (Fig. 1A-C). Given previous studies showing different population dynamics across cortical layers (Orduz et al., 2019), we analysed the upper (Bin 1) and lower (Bin 2) parts of the cortex separately (Fig. S1). In the cortex, Sox9+ cells are predominantly glial progenitors at P1 and P5 and astrocytes from P15 (Fig. 1A). No difference was observed in Sox9+ cell density between the upper and lower cortical parts at any stage (Fig. 1B, D and Table S1). Density peaked at P5, with a significant increase between P1 and P5 followed by a significant decrease until P60. This decrease was particularly pronounced between P5 and P15 (Bin 1: 2.4-fold; P5: 1857±137; P15: 780±70.49; P60: 426.7±69.89 10^2^/mm^3^ cells; Bin 2: 2.3-fold; P5: 1816±181; P15: 775±34.66; P60: 359.7±110.8 10^2^/mm^3^ cells). Sox10+ labelling was too faint to be analysed at P1. After P5, a small reduction in Sox10+ cell density was also observed in Bin 1 and after P15 in Bin 2 (Fig. 1C,E). A higher cell density was found in the lower part of the cortex (Bin 2), at all studied stages (Fig. 1C,E and Table S1). We then compared the two cell populations. Sox9+ cells were more than 2 times more abundant than Sox10+ cells in the upper part of the cortex at P5 (Fig. 1B-E and Table S1). In the lower part, Sox9+ cells were more abundant at P5, but Sox10+ density was higher at P15 and P60 (Fig. 1B-D and Table S1). Altogether, we reveal disparities in density between Sox9+ and Sox10+ cell populations during cortical postnatal development.

**Figure 1.**
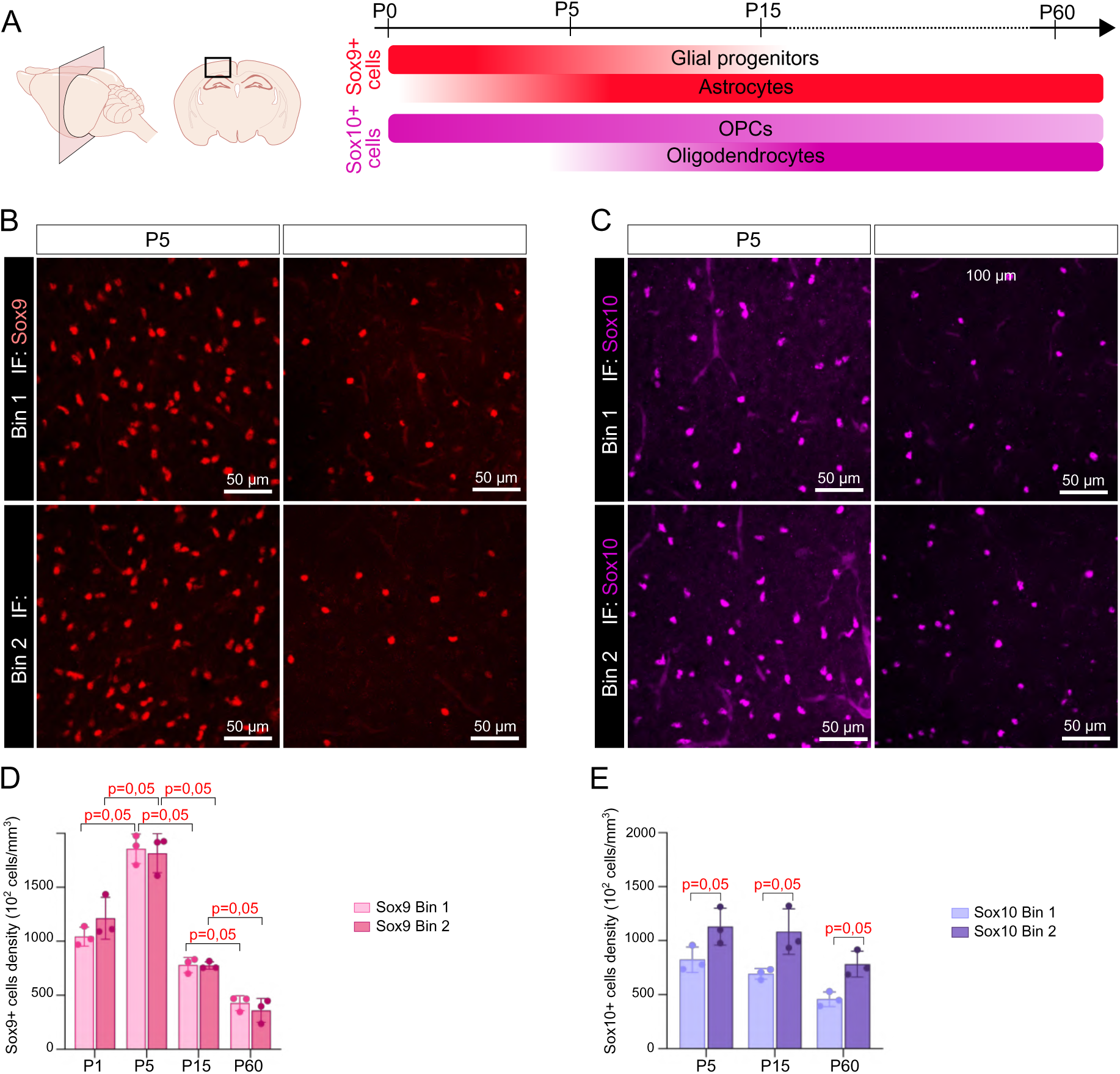
Macroglial cells density during postnatal cortical development. **A.** Schematic representation of the studied area in the mouse brain (left) and developmental timeline (right) of the macroglial cells expressing Sox9 (red) or Sox10 (magenta). **B, C.** Spinning disk W1 Z-stack projection images of Sox9+ (red) (B) or Sox10+ (magenta) (C) cells detected by immunofluorescence at P5 and P60 in Bin 1 or Bin 2 of somatosensory cortical sections. **D, E.** Analysis of the Sox9+ (D) and Sox10+ (E) cells densities shown as a number of cells x10^2^ per mm^3^. N= 3 mice, n=1 section per animal, results are shown as mean of the results on 4 ROI ± SD. Statistics: Mann Whitney test between ages (D) or between layers (E).

### Distribution of macroglial cells in the cerebral cortex during postnatal development

We next analysed the distribution of macroglial cells during postnatal development of the somato-sensory cortex. First, we measured the mean distance between each cell and its nearest neighbour. Consistent with the decreased cell density of Sox9+ cells after P5, the mean distance between neighbouring Sox9+ cells progressively increased after P5 (Bin 1; P5: 12.6±0.2; P15: 17.1±0.2; P60: 22.5±1.4 µm) (Bin 2; P5: 13.0±0.5; P15: 18.0±0.5; P60: 25.6±0.9 µm). At P60, Sox9+ cells were also slightly closer to each other in the upper compared to the lower part of the cortex (Fig. 2A). Measurement of the mean distance between each Sox9+ cell and its 10 nearest Sox9+ cells gave similar results (Fig. 2B, Table S1). For Sox10+ cells, no difference in the mean distance between adjacent cells nor with 10 nearest cells was observed between P5 and P60 (Fig. 2C, D and Table S1). Consistent with density differences (Fig. 1), Sox10+ cells were closer to each other in the lower part of the cortex from P5 (Fig. 2C, D and Table S1).

**Figure 2.**
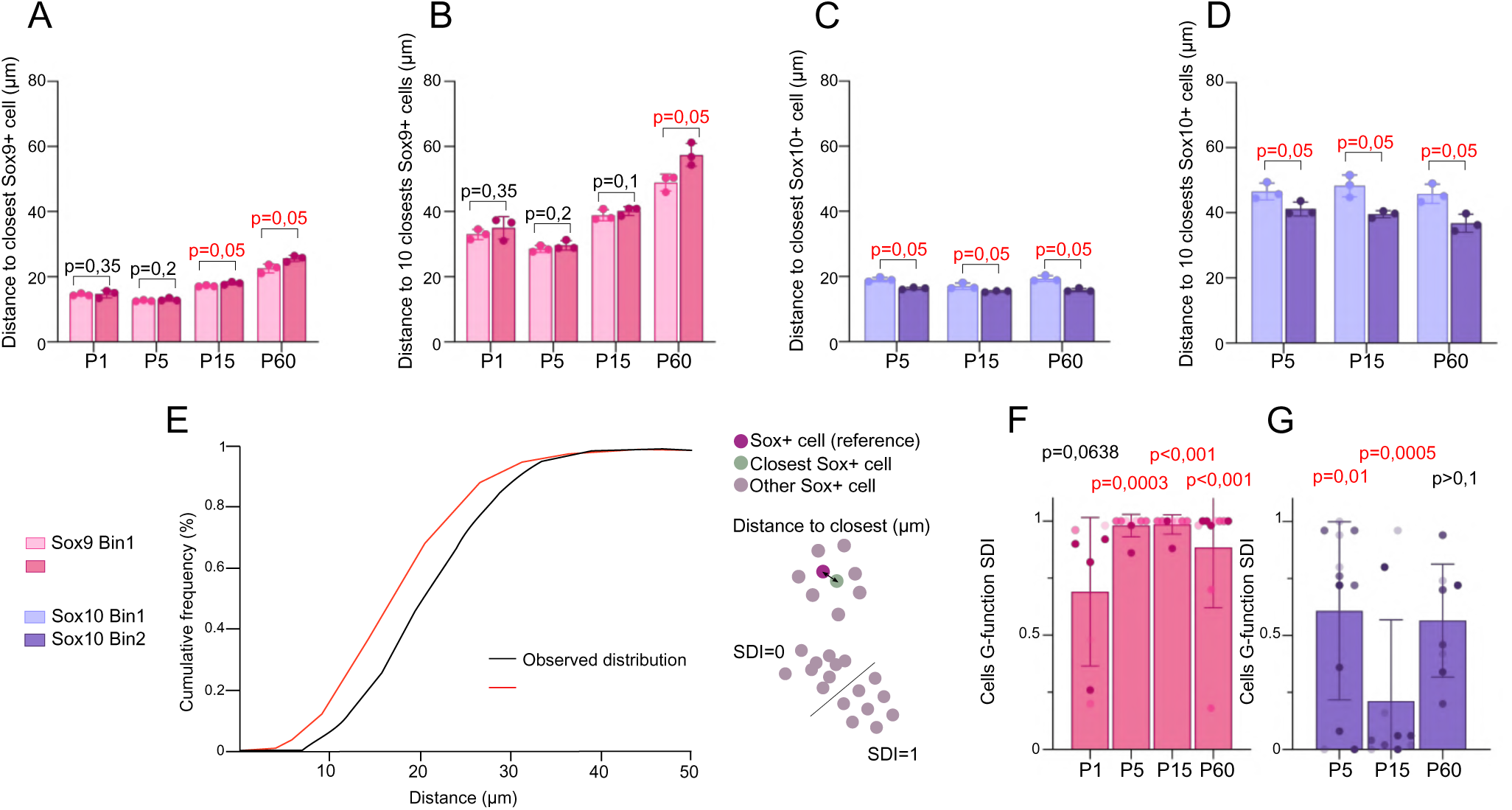
Macroglial cells distribution during postnatal cortical development. **A, B.** Analysis of mean distance to the closest (A) or 10 closest (B) Sox9+ cells, shown as the mean distance for each cell of the studied area in the somato-sensory cortex. **C, D.** Analysis of mean distance to the closest (A) or 10 closest (B) Sox10+ cells, shown as the mean distance for each cell of the studied area in the somato-sensory cortex. **E**. Schematic summary of the G-function analysis of cell distribution. Cells can follow three main behaviours: grouped (SDI=0), randomly distributed or regularly spaced (SDI=1). The G-function is the cumulative distribution function (CDF) of the distance between a typical point (the labelled cell) in the pattern and it nearest-neighbour. The experimental CDF (black curve) is obtained for each ROI analysed and is compared to the CDF of randomly generated point patterns in the same area and with the same number of points (red curve). For each ROI analysed, comparison between the experimental and random CDF are computed as the Spatial Distribution Index (SDI). Distribution of SDIs in each experimental condition indicates the distribution pattern followed by the cells. **F, G**. Result of G-function analysis for Bin 2 Sox9+ (F, pink) and Sox10+ (G, purple) cells across postnatal development. N= 3 mice, n=1 section per animal, results are shown as mean of the results on 4 ROI (A-D) per section or the result of one ROI (F) ± SD. Statistics: Mann Whitney test between ages (A,B) or layers (C,D). F, G Kolmogorov-Smirnov test.

The spatial arrangement of cells within a population can vary from clustered to evenly spaced patterns, which may reflect distinct developmental processes such as cellular attraction or repulsion. To analyse this distribution for macroglial cells across development, we adapted a the *Fiji* plugin “*2D/3D Spatial Analysis”* to assess the deviation from spatial randomness (Andrey et al., 2010). We computed the G-function, the cumulative distribution function (CDF) of the distance between a labelled cell and its nearest-neighbour. Shorter distances suggest clustering, while longer distances indicate regular spacing (Fig. 2E). The experimental CDF was compared to the CDF of randomly generated patterns within the same area and with the same number of cells. For each region of interest (ROI), this comparison was computed as the Spatial Distribution Index (SDI) (Andrey et al., 2010). Distribution of SDIs indicates the cell distribution pattern: uniformly distributed SDIs from 0 to 1 reflects a random distribution of cells. SDIs near 0 suggest clustering. SDIs near 1 indicate regular spacing (Andrey et al., 2010) (Fig. 2E).

Given the higher density of Sox10+ cells in the lower part of the cortex, we first focused our study on this region. For Sox9+ cells, the SDIs distribution was significantly different from the random distribution from P5 (Fig. 2F, Table S1), and values were not different from 1 (Bin 2: P5: 0.98±0.05; P15: 0.98±0.04; P60: 0.88±0.3) (Wilcoxon signed Rank Test: P5, p=0.5; P15, p>0.9, P60, p=0.1), revealing a regular distribution. At P1, however, SDIs were not different from normal distribution and not different from 1, indicating a more random cell distribution (Wilcoxon signed Rank Test: P1: p=0.008). Same results were found in the upper layers of the cortex (Table S1). These results indicated that, from P5 onwards, Sox9+ cells are regularly organized.

For Sox10+ cells, the SDIs distribution differed between stages. At P5 and P15, distribution was significantly different from random but not at P60 (Fig. 2G; Table S1). However, SDIs were always significantly different from 1 (Wilcoxon signed Rank Test: P5, p=0.001; P15, p=002, P60, p=0.008). The same result was found in the upper part of the cortex (Table S1). This indicates that organisation of Sox10+ population always differs from a regular distribution and the mature random distribution is established by P60.

Overall, our findings demonstrate that Sox9+ and Sox10+ cells follow distinct postnatal trajectories adopting specific distribution patterns in the somatosensory cortex. Sox9+ cells maintain a regular distribution from P5; Sox10+ cells are denser in deeper cortical layers, are never regularly organized and change their spatial distribution during postnatal development.

### Distribution of glial cells relative to the vascular compartment

Macroglial cells are closely associated with the brain vasculature. In the adult CNS, astrocytes entirely cover the BVs through specialized processes called endfeet or perivascular astrocyte processes (PvAP) (Cohen-Salmon et al., 2020). The formation of PvAPs, which occurs from birth to P15, plays a crucial role in BV maturation (Cohen-Salmon et al., 2025; Freitas-Andrade et al., 2023; Gilbert et al., 2021, 2019; Mondo et al., 2020). Furthermore, parts of OPCs migrate along BVs during embryonic development (Lepiemme et al., 2022).

To better understand the relationship between macroglial cells and BVs during postnatal development, we immunolabeled for Pecam-1, an endothelial cell-specific protein (Fig. 3A, B). The cortical vascular volume, normalized by the cortical volume, changed during postnatal development: while stable between P1 and P5, a global increase in vascular volume was observed between P5 and P15 (Fig. 3C) due to postnatal angiogenesis and capillary bed extension (Coelho-Santos and Shih, 2020) (Bin 1; P1: 0.08±0.009; P5: 0.1±0.01; P15: 0.2±0.03 µm^3^/µm^3^) (Bin 2; P1: 0.08±0.007; P5: 0.08±0.01; P15: 0.1±0.02 µm^3^/µm^3^). After P15, the vascular volume decreased, probably related to the cortical growth (Fig. 3A, D and Table S1). Vessel branching followed the same trends (Fig. 3F and Table S1). Of note, the vascular volume and branching were globally slightly reduced in the lower part of the cortex compared to the upper part but this difference was only significant at P15 for the branching (Fig. 3D, E, F and Table S1).

**Figure 3.**
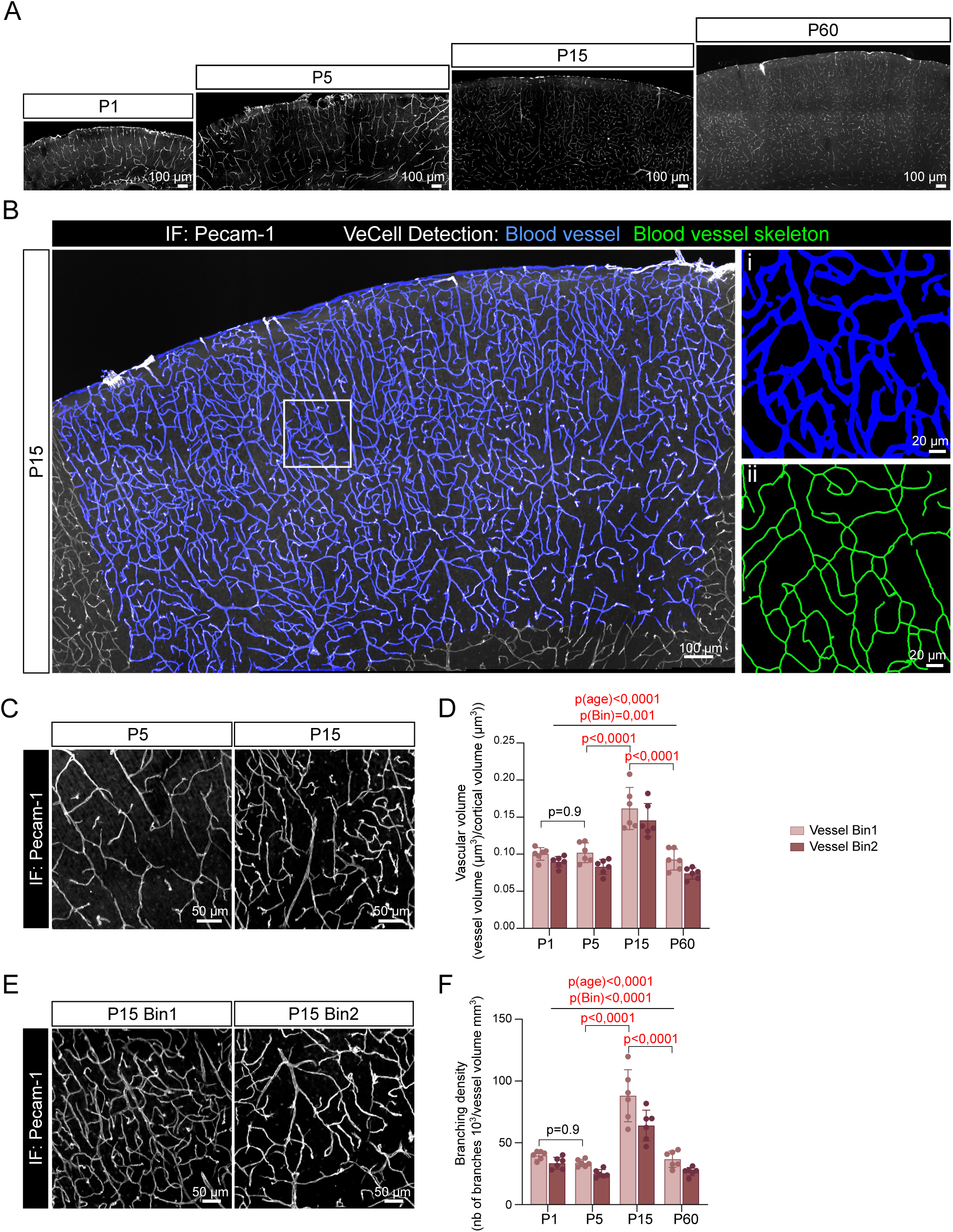
Vascular system organisation during postnatal cortical development. **A.** Representative Axio Zoom microscopy image of P1, P5, P15 and P60 somatosensory cortical section stained for blood vessels (white) detected by immunofluorescence on Pecam-1. **B**. Representative Axio Zoom microscopy Z stack projection image of P15 somatosensory cortical section stained for blood vessels (white) detected by immunofluorescence on Pecam-1 with the superimposed masks created by the VeCell Fiji plugin (in blue). i and ii are detailed of boxed area. i, BVs mask created by the VeCell Fiji plugin (blue); ii, skeleton mask of the BV for branching analysis (green). **C**. Representative Axio Zoom microscopy Z-stack projection images of P5 and P15 somatosensory cortical sections in Bin1, stained for blood vessels detected by immunofluorescence for Pecam-1 (white). **D.** Analysis of the vascular volume reported as BVs volume normalized by cortical volume. **E**. Representative Axio Zoom microscopy Z-stack projection images of P15 somatosensory cortical sections Bin1 (left) or Bin2 (right), stained for blood vessels detected by immunofluorescence for Pecam-1 (white). **F**. Analysis of the vascular branching as the number of 10^3^ branches per mm^3^ of BVs. N= 3 mice, n=2 sections per animal. Results are shown as mean of the results on 4 ROI per section ± SD. Statistics: 2-way ANOVA; p(age): result of ANOVA for age effect; p(Bin): result of ANOVA for cortical layer effect.

We next analysed the distance between macroglial cells and their nearest BV in the somatosensory cortex at all stages (Fig. 4.A). We calculated both the mean distance to BVs for all cells (Fig. 4C and E) and the percentage of cells with the nucleus directly apposed to BVs (Fig. 4D, F). The distance to BV was the highest at P5, and similar for Sox9+ and Sox10+ cells (Fig. C-E, Table S1). In contrast, from P15, Sox9+ cells were closer to BVs than Sox10+ cells and a higher proportion of Sox9+ were found apposed to BV (Fig. 4B-D, Table S1).

**Figure 4.**
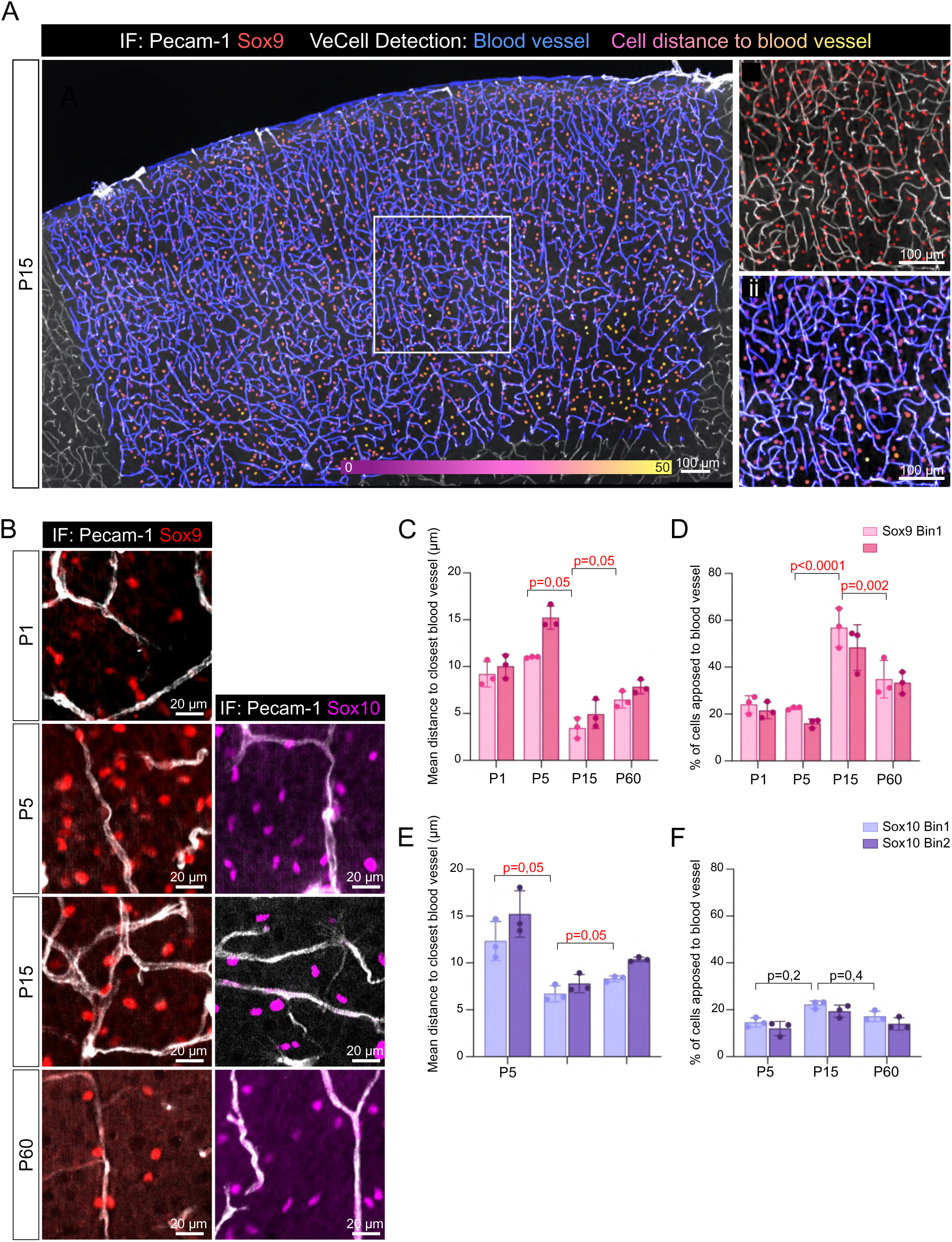
Macroglial cells distribution in relation to blood vessels during postnatal cortical development. **A.** Representative Axio Zoom microscopy Z-stack projection image of a P15 somatosensory cortical section stained for blood vessels (white) detected by immunofluorescence on Pecam-1 with the superimposed masks created by the VeCell Fiji plugin. BVs mask is in blue and Sox9+ cells are color coded for their distance to nearest BV from magenta (close to BV) to yellow (far from BV). **i and ii** are detailed of boxed area. i, immunofluorescence of Pecam-1-labeled BVs (white) and Sox9+ cells (red); ii, BVs mask created by the VeCell Fiji plugin (blue) superimposed to the Pecam-1 staining (white) and mask of Sox9+ cells color-coded for their distance to nearest BV. **B.** Representative Axio Zoom microscopy Z-stack images of Sox9+ (red) or Sox10+ (magenta) cells and Pecam-1 (white) detected by immunofluorescence at P1, P5, P15 and P60 in Bin 2 of somatosensory cortical sections. **C, E.** Analysis of the mean distance to nearest BV of Sox9+ (C) and Sox10+ (E) cells. **D, F.** Analysis of the Sox9+ (D) and Sox10+ (F) percentage of nuclei apposed to BV (distance to nearest BV=0 µm). N= 3 mice, n=1 section per animal. Results are shown as mean of the results on 4 ROI per section ± SD. Statistics: Mann Whitney test between ages (C, E) and chi-square test between ages (D, F)

Altogether, we show here that macroglial cells are more distant to BVs at P5. From P15, Sox9+ cells are closer to BVs than Sox10+ cells, suggesting distinct interactions between these macroglial populations and the vasculature. Thus, we reveal different developmental changes in the spatial relationship between macroglial cells and BVs.

### Macroglial cell density and distribution are not altered in *Mlc1* KO cortex

Whether the organisation of macroglial cells can be altered in pathological contexts remains poorly studied. MLC1 is a membrane protein expressed by the astrocyte lineage in the brain whose absence leads to Megalencephalic Leukoencephalopathy with subcortical Cysts (MLC), a rare type of leukodystrophy (Passchier et al., 2024). MLC1 expression starts at P5. The protein progressively localizes within astrocyte perivascular processes (PvAP) and forms a junctional complex between PvAPs with GlialCAM, which is mature around P15 (Gilbert et al., 2019; Teijido et al., 2007). In *Mlc1* KO mice, the gliovascular interface is altered during development: astrocyte polarity and vascular coverage do not develop properly and vascular functions are altered, including arteriolar contractility, CSF drainage and neurovascular coupling (Gilbert et al., 2021). Thus, we hypothesized that absence of MLC1 could also perturb astrocyte spatial arrangement.

On P60 somatosensory cortical sections, we observed no difference in the vascular volume and branching (Fig. 5A-D). We found no difference in Sox9+ cell density (Fig. 5A and E). We observed however a slight decrease in the distance between cells in *Mlc1* KO (Bin 1; *Mlc1* WT: 23.4±1.7; *Mlc1* KO: 20.6±1.8 µm; Bin 2; *Mlc1* WT: 27.6±1.2; *Mlc1* KO: 23.7±1.6 µm). G-function-assessed distribution pattern was similar between genotypes (Fig. 5H) and similar to our previous observation for Sox9+ cells: SDI distribution different from the normal distribution and not different from 1 (Fig. 5G and Fig. 2). Finally, the mean distance of Sox9+ cells to the closest BV and the percentage of Sox9+ cells apposed to blood vessels were similar in *Mlc1* KO and controls (Fig. 5I, J). Mlc1 is expressed in astrocytes as early as P5 and we previously identified a transient alteration of perivascular coverage in the *Mlc1* KO mouse cortex (Gilbert et al., 2021, 2019). To determine whether this effect extends to astrocyte distribution and density we analysed *Mlc1* KO at P5 and P15. We confirmed the absence of difference in the vascular organisation (Fig. S2). As to astrocytes, we found no differences in astrocyte density, distribution nor distance to blood vessels at both P5 and P15 (Fig. S2).

**Figure 5.**
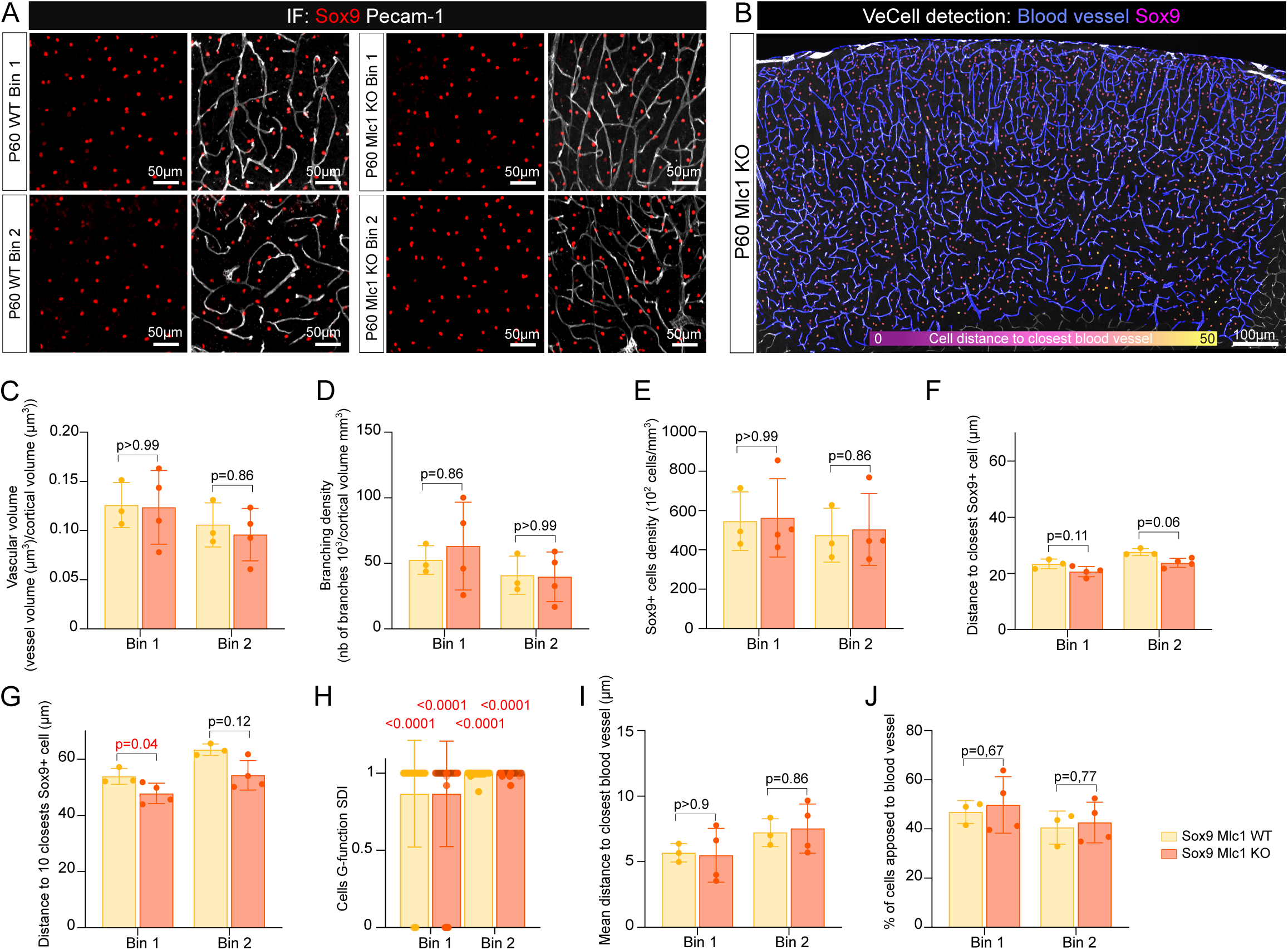
Sox9+ cell density and distribution in the *Mlc1* KO model. **A.** Spinning disk W1 Z-stack projection images of P60 upper and lower parts of the somatosensory cortex after immunostaining for Sox9 (red) and BV (grey). **B**. Masks created by the VeCell Fiji plugin on P60 *Mlc1* KO Axio Zoom microscopy Z-stack projection image, BVs mask is in blue and Sox9+ cells are color coded for their distance to nearest BV from magenta (close to BV) to yellow (far from BV). **C.** Analysis of the vascular volume reported as BVs volume normalized by cortical volume. **D.** Analysis of the vascular branching as the number of 10^3^ branches per mm^3^ of BVs. **E**. Analysis of the Sox9+ cells densities shown as a number of cells x10^2^ per mm^3^. **F, G**. Analysis of mean distance to the closest (F) or 10 closest (G) Sox9+ cells, shown as the mean distance for each cell of the studied area in the somato-sensory cortex. **H**. Result of G-function analysis for Sox9+ cells in Bin 1 and 2 in the two genotypes. **I.** Analysis of the mean distance to nearest BV of Sox9+ cell. **J.** Analysis of the Sox9+ percentage of nuclei apposed to BV (distance to nearest BV=0 µm). N= 3 WT and 4 *Mlc1* KO mice, n=3 section per animal, results are shown as mean of the results on 4 ROI per section ± SD. Statistics: C,D,E,F,G,I: Mann-Whitney test between genotypes, H: Kolmogorov-Smirnov test, J: Chi square test

We thus show that the vascular network as well as the distribution and density of Sox9+ cells, remain unaffected in the absence of MLC1.

## Discussion

Using a newly developed Fiji plugin, we revealed distinct distribution postnatal developmental patterns for astrocytes and oligodendrocytes. The density of Sox9+ cells was highest at P5 compared to P1 and P15/P60, similar to observations using Aldh1l1 as astrocyte-specific marker (Freitas-Andrade et al., 2023). We showed a higher density of Sox9+ cells compared to Sox10+ at P5 but not at P60, consistent with the reported intense proliferation of astrocyte progenitors around P4 (Clavreul et al., 2019) and the later reduced expression of Sox9 in progenitors differentiating into OligoLC (Klum et al., 2018). We observed a smaller Sox10+ population in the upper part of the somatosensory cortex at all developmental stages, as previously described for OPCs (Ogino et al., 2020). This effect has been shown to rely on Reelin secreted by Cajal-retzius cells during late embryonic development, which acts as a driving force for OPC migration (Barber et al., 2022; Ogino et al., 2020). It is also consistent with the known higher density of myelinating oligodendrocytes in the lower cortical layers (Orduz et al., 2019). We observed a random distribution of Sox10+ cells within cortical layers as previously showed (Orduz et al., 2019). In contrast, we showed that Sox9+ density was identical between layers of the cortex and displayed a regular distribution indicating similar distances between cells. This is in line with the well-established organisation of astrocytes which occupy exclusive, non-overlapping domains that completely tile the cortical parenchyma in the mature brain, with each cell occupying a comparable volume (Bushong et al., 2004, 2002; Halassa et al., 2007). We observed an increase of the vascular volume and branching between P5 and P15, in line with the postnatal angiogenesis phase between P5 and P15 (Coelho-Santos and Shih, 2020; Freitas-Andrade et al., 2023).

A key aspect of our study was the description of distinct patterns of distribution for macroglial cells. We found Sox9+ astrocytes to be regularly distributed within the parenchyma as early as P5, suggesting an extremely precise and coordinated colonization of the entire cortical parenchyma by macroglial progenitors/astrocytes. In contrast, Sox10+ cells were more randomly distributed. Astrocyte tiling of the parenchyma is crucial to neuronal function (Baldwin et al., 2021). Such very early acquired organisation suggests the presence of active mechanisms of active attraction and repulsion mechanisms between astrocyte progenitors which remain to be fully understood. In contrast, Sox10+ cells displayed a random distribution, suggesting different mechanisms orchestrating oligodendrocytes and OPCs postnatal colonization of the cortical parenchyma.

The interaction between macroglial cells and BVs has gathered increasing interest in both developmental and disease contexts (Lepiemme et al., 2022; Niu et al., 2019; Watkins et al., 2014). After P5, both Sox9+ and Sox10+ cells were found to be closer to BVs. This result might simply reflect the increased BVs density in the parenchyma. However, active migration mechanisms driven by vascular signals to attract astrocytes could also be involved. BVs have been shown to participate in the maturation of astrocyte perivascular processes (Cohen-Salmon et al., 2025). Some of these mechanisms may also promote astrocyte attraction, thereby contributing to their distribution and regular spacing, including oxygen level (Uemura et al., 2006; West et al., 2005), secreted growth factors such as Pdgfβ (Daneman et al., 2010; Munk et al., 2019), and extracellular matrix proteins such as laminin (Menezes et al., 2014; Segarra et al., 2018; Yao et al., 2014) or Fibronectin (Liu et al., 2023).

Interestingly, Sox9+ nuclei were closer to BVs than Sox10+ which is consistent with the known higher interaction between astrocytes and BVs, compared to oligodendrocytes, including the complete coverage of BVs with PvAPs by P15 (Gilbert et al., 2021; Mathiisen et al., 2010).

In order to take our study a step further, we analyzed a mouse model deficient for the astrocyte protein MLC1, in which astrocyte polarity and PvAP formation are disrupted (Gilbert et al., 2021). We found no difference in vascular organisation, consistent with our previous report (Gilbert et al., 2021). The regular astrocyte spatial distribution was also preserved. Thus, the altered perivascular coverage in this model does not imply a change in astrocyte distribution. We cannot however exclude the possibility of transient developmental changes as previously reported in this model (Gilbert et al., 2021). Defects in macroglial production and/or distribution could be further analyzed in various cerebrovascular and neurodevelopmental disorders that emerge during late brain development, such as cavernomas, autoimmune diseases including multiple sclerosis and Neuromyelitis Optica, arteriovenous malformations, muscular dystrophies affecting the brain, and other leukodystrophies (Cohen-Salmon et al., 2025). While the destruction or abnormal proliferation of astrocytes, oligodendrocytes, and OPCs have been frequently documented in diseases of the mature brain, impairments in their production and spatial organization during postnatal development—whether or not related to the vascular system—remain largely unexplored.

Altogether, we analyzed macroglia organization during postnatal development using a reliable, freely available Fiji plugin using simple brain sections and low-resolution microscopy. This tool successfully reproduces previously reported findings, highlighting the robustness of our approach. We revealed strong differences between the developmental trajectories of astrocyte and oligodendrocyte lineages. Notably, we observed that astrocytes adopt a remarkably regular distribution pattern that is established very early during postnatal development.

### Limitations

Most of the data presented in this article were obtained form one or two sections per mouse (n=3). In a preliminary study conducted at P15, no differences were observed between data obtained from a single section with the mean value calculated from four sections. However, we cannot exclude limited accuracy in the assessment of spatial distribution and blood vessel architecture, as these parameters may be influenced by sample preparation.

## Acknowledgments

We gratefully acknowledge the Orion technological core (IMACHEM-IBiSA) of Center for Interdisciplinary Research in Biology for their support, and especially Estelle Anceaume and Julien Dumont for assisting us with the Axio Zoom microscopy image acquisition. We thank the whole “physiology and physiopathology of the gliovascular unit” team for helpful comments and contribution on the VeCell plugin functionality. We would like to thank, in particular, Camille Claveau and Barbara Delaunay for their help with the study of *Mlc1* KO mice.

## Author contributions

Conceptualization, NG and A-CB; Methodology, NG, HM, PM and A-CB; Investigation, NG and TH; Writing-Original Draft, A-CB; Writing-correction, NG, MC-S and A-CB; Funding Acquisition, MC-S. and A-CB; Supervision, A-CB.

## Data Availability

The data that support the findings of this study are available from the corresponding author upon reasonable request.

---

**Supplementary Figure 1.**
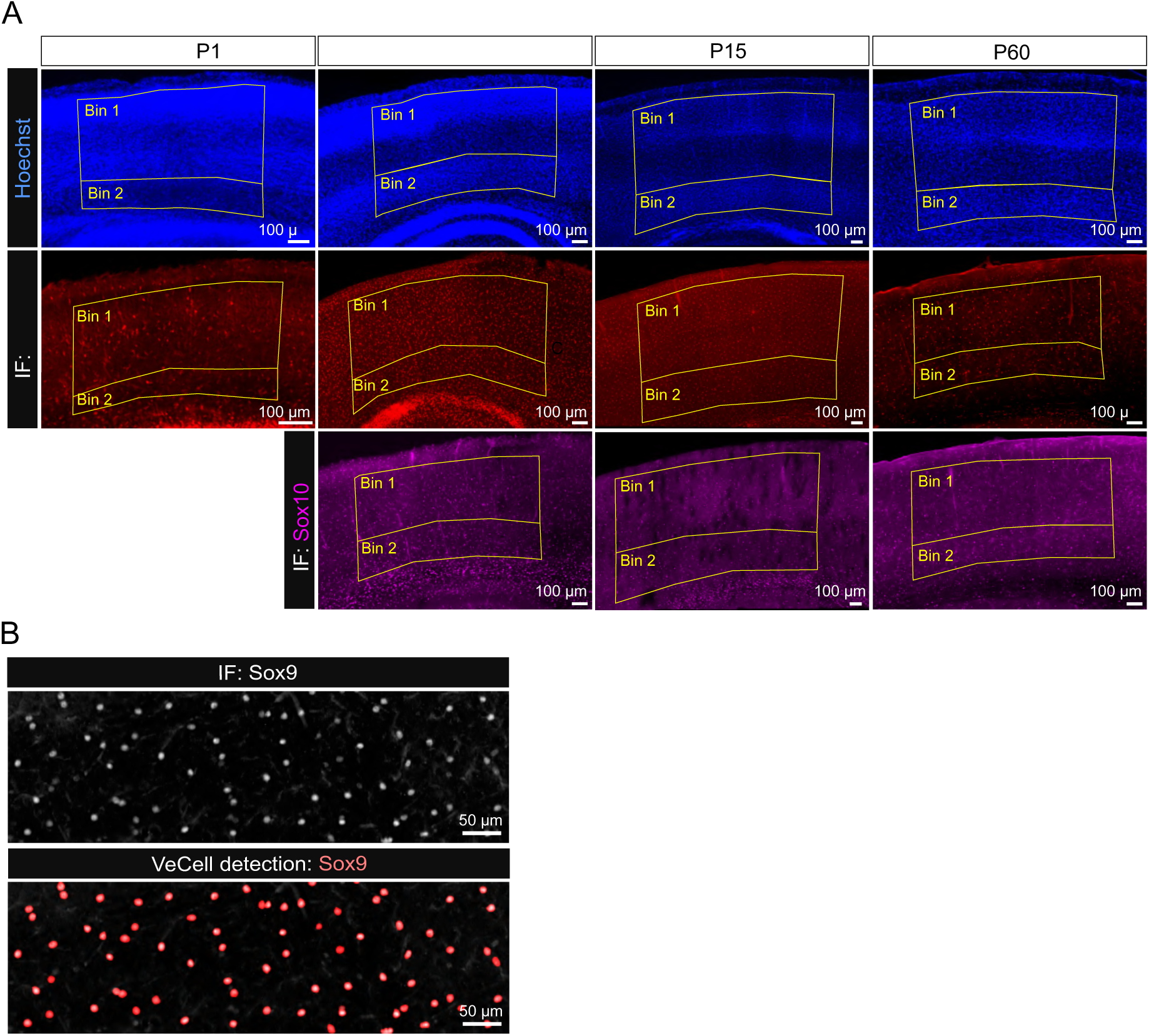
**A**. Representative Axio Zoom microscopy images of Hoechst (blue), Sox9+ (red) or Sox10+ (magenta) showing the Region Of Interest within the somato-sensory cortex (Bin 1: upper layers (II/III-V), Bin 2: lower layer (VI)). **B**. Representative Axio Zoom microscopy Z-stack projection image of Sox9+ cells detected by immunofluorescence (top, white) on a P15 somatosensory cortical slice and the mask created by the *VeCell* Fiji plugin (down, red).

**Supplementary Figure 2.**
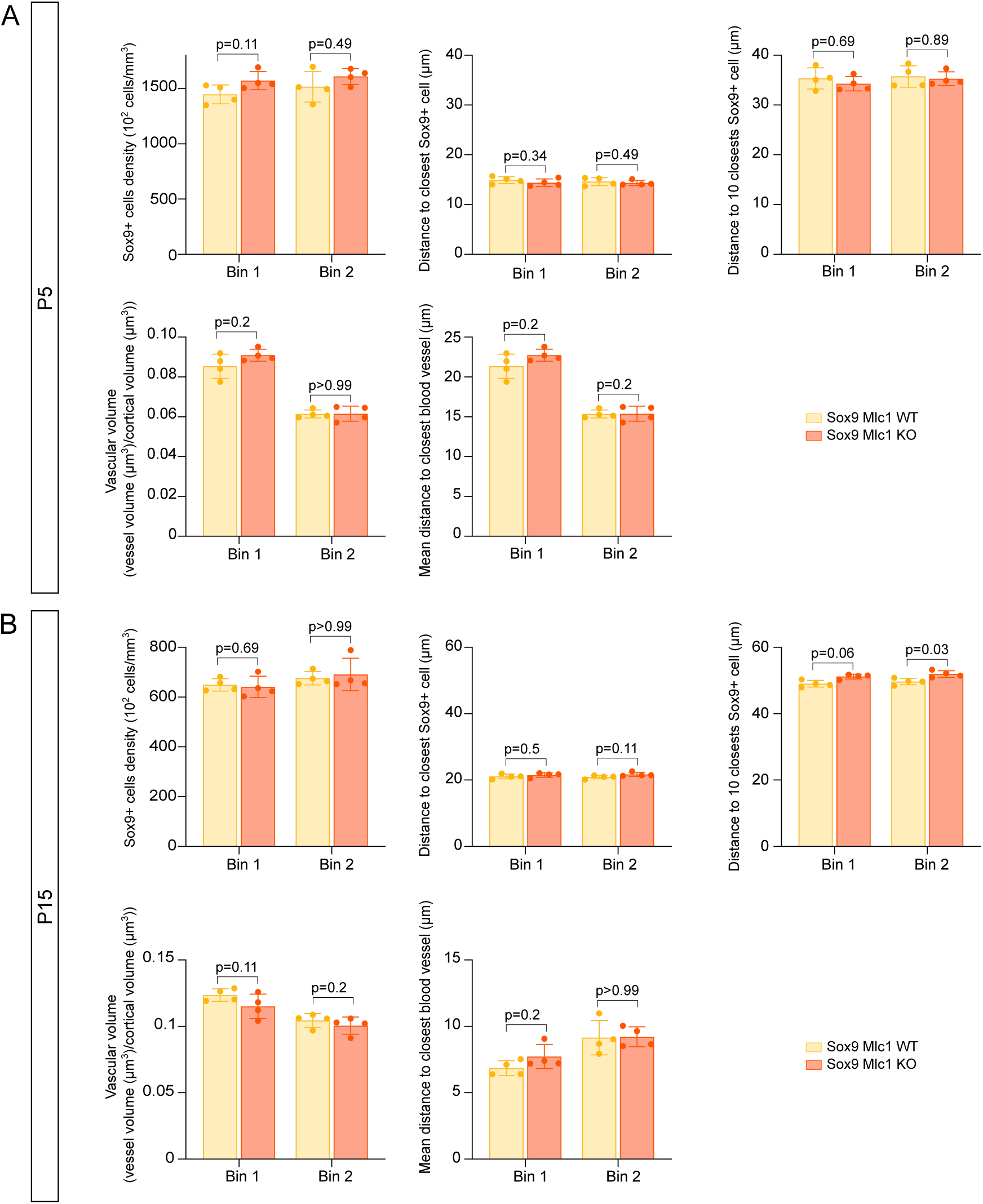
Sox9+ cell density and distribution during development in the *Mlc1* KO model. Analyses at P5 (A) and P15 (B) of Sox9+ cells density shown as a number of cells x10^2^ per mm3, mean distance to the closest or 10 closest Sox9+ cells, shown as the mean distance for each cell of the studied area in the somato-sensory cortex, vascular volume reported as BVs volume normalized by cortical volume, mean distance to the closest BVs in µm. N= 4 mice, n=3 sections per animal. Results are shown as mean of the results on 4 ROI per section ± SD. Statistics: Mann Whitney test between genotypes.

**Supplementary Table 1** List of data and results of statistical tests used in this article

